# A systematic approach to understanding how patient variants affect the activity of Wiskott-Aldrich syndrome protein

**DOI:** 10.64898/2026.05.19.726146

**Authors:** Lia G Pinho, Marcus R Bezerra, Axel Leppert, Rhaissa C Vieira, Katharina Riffelsberger, Julia Schwekendiek, Minghui He, Marton Keszei, Julien Record, Ulf Tedgård, Fredrik Kahn, Mikael Sundin, Annick van de Ven, Jolanta Bernatoniene, Kim Ramme, Alejandro Palma, Anna Shcherbina, Olov Ekwall, Gilvan Pessoa Furtado, Michael Landreh, Lisa S Westerberg

**Author notes:** **Corresponding Authors: Lia G Pinho,** Ph.D., Karolinska Institutet, Department of Microbiology, Tumor and Cell biology, Biomedicum C7, Stockholm, Sweden, **Lisa S. Westerberg**, Ph.D., Karolinska Institutet, Department of Microbiology, Tumor and Cell biology, Biomedicum C7, Stockholm, Sweden.

## Abstract

Wiskott-Aldrich syndrome (WAS) and X-linked neutropenia (XLN) are caused by genetic variants in the *WAS* gene. How *WAS* variants lead to clinical disease remains unsolved in many cases. We expressed human WASp using a spider silk inspired solubility tag (NT*-tag) and inserted patient’s variants. Native mass spectrometry and pyrene actin assays showed that five variants (L270P, F271S, S272P, I290T, I294T) predicted to cause XLN led to open protein conformation and high actin polymerization rate in the absence of the WASp activator, Cdc42. One previously reported XLN variant (R268W), two loss-of-function WAS variants (A236G, D485N), and one variant of unknown significance (R431W) behaved similarly to wildtype WASp in terms of structural conformation and actin polymerization. Patient CD4^+^ T cells were used for analysis of WASp expression and phosphorylation, actin polymerization, anti-CD3 induced proliferation capacity, and upregulation of high affinity LFA-1, distinguishing loss-of-function and gain-of-function variants from benign *WAS* variants. This systematic approach reveals how *WAS* genetic variants cause severe human disease and stratify variants to guide clinical decision for definitive therapy.

**Key Points:**

- Gain-of-function WASp variant has extended protein conformation probed by native mass spectrometry and raised pyrene actin polymerization.
- Functional analysis of patients CD4^+^ T cells classifies WASp variants as loss-of-function, reduced-function, gain-of-function, and benign.

## Introduction

Actin cytoskeleton is essential for hematopoietic cells in regulating cell movement, cell–cell interactions, cell signaling, and cell division. Genetic variants in actin-related proteins can alter this dynamic process, as observed in immunoactinopathies^1^. Wiskott-Aldrich syndrome (WAS), caused by the loss-of-function (LOF) variants in the *WAS* gene encoding actin nucleation promoting factor (NPF) WAS protein (WASp). WAS patients suffer from a triad of immunodeficiency, eczema, and thrombocytopenia and have increased risk of developing autoimmune disease and hematological malignancies ^2,3^. WAS patients can be classified into two groups based on the *WAS* gene variants. Class I variants include missense variants in exons 1-2 and the intronic hot spot variant c.559+5G>A, typically associate with residual WASp expression and milder disease. Class II variants include all other variants, often leading to absent WASp expression and early onset of severe disease^4^. In contrast to WAS, X-linked neutropenia (XLN) is caused by predicted gain-of-function (GOF) variants in the *WAS* gene. It is challenging to determine if a new WAS variant of unknown significance (VUS) is disease causing and a standardized set of assays is lacking. Diagnosis of WAS and disease prognosis are determined by combining disease manifestations, class of genetic variant, and WASp expression measured by flow cytometry. For XLN in which WASp expression is normal, only the *WAS* gene variant and congenital neutropenia are used to establish diagnosis. However, neutropenia in children can be caused by other reasons, and functional assay for suspected XLN patients is lacking.

WASp is expressed uniquely in hematopoietic cells^5,6^. It is a highly disordered protein that has been difficult to express in full length, preventing the biochemical analysis to determine how *WAS* variants affect protein function. WASp assumes an autoinhibited conformation by intramolecular interaction between the GTPase-binding domain (GBD) and the verprolin cofilin acidic (VCA) domain^7–9^. Cdc42-GTP binding to the GBD disrupts the hydrophobic core and release the VCA, enabling its interaction with G-actin and the Arp2/3 complex ^7–9^ to induce branched actin polymerization. XLN, GOF, variants located in the GBD are predicted to destroy autoinhibition and constitutively expose the VCA domain. WASp contains an N-terminal WASp homology (WH)1 domain that interacts with WASp interacting protein (WIP), that prevents proteasomal degradation of WASp by inhibiting WASp ubiquitylation ^10,11^. Release of the autoinhibition exposes WASp-tyrosine(Y)291 in the GBD domain that when phosphorylated, sustains WASp activity toward actin polymerization by Arp2/3, even upon dissociation from Cdc42^12^.

A systematic functional comparison of *WAS* class I and class II, XLN, and VUS variants is lacking. We stably expressed NT*-tagged WASp and performed native mass spectrometry to probe conformational states and pyrene actin polymerization assays. We set up functional assays using CD4^+^ T cells for prediction of WAS and XLN patient variants. Comparative characterization of 11 *WAS* variants demonstrated that these assays can be used to identify molecular and cellular effects of WASp function and help guiding treatment of WAS and XLN patients with new *WAS* variants.

## Methods

### Patients

Patients, their family members, and unrelated healthy controls that were sex and age matched were enrolled. We studied 9 suspected WAS patients and 3 XLN patients (previously reported^13^). The Code of Ethics of the World Medical Association (Declaration of Helsinki) for human samples was followed; the study was approved by the institutional Ethical Committee (EPM# 2023-04773). The family caregivers and patients, where possible, have provided written consent.

### Structural analysis

Alphafold 3.0^14^ was used to obtain WASp structure to display the sites of mutations. Figure was made by PyMOL molecular viewer (https://www.pymol.org/). The potential effect of the variants was predicted using DDmut^15^.

### Native Mass Spectrometry

Mass spectra were recorded using a Waters Synapt G1 adapted for the analysis of intact protein complexes (MS Vision, Netherlands) and equipped with an offline nanospray ionization source. The proteins were desalted using ZebaSpin (7k) columns (Thermo Scientific) and loaded into ESI capillaries (Thermo Scientific®). The instrument operated at a capillary voltage of 1.5 kV, cone voltage of 20 V and extraction cone voltage of 4 V. The source temperature was set to 30°C while the source pressure was kept at 8 mbar. As the trap gas, argon was used at a flow rate of 4 mL/min. The collision energy in the ion trap was set to 20 V and the transfer of collision energy at 10 V. To visualize the acquired mass spectra, Mass Lynx 4.1 (Waters, UK) was used, while the weighted average charge was calculated in mMass (v3.9.0.).

### Actin Polymerization Assay

Pyrene actin was used to follow actin polymerization as described previously^16^. G-actin (2 μM, 10% pyrene-labeled; Cystoskeleton, Inc) was mixed with 10 nM Arp2/3 complex (Hypermol®) and 1µM WASP in KMEI buffer. The polymerization was recorded in the SpectraMax iD3 (Molecular Devices) at an excitation wavelength of 365 nm and emitted fluorescence at 407 nm. The measurement interval was set at 20 seconds at a medium gain with a reading duration of 60 minutes.

### Analysis of WASp F-actin content by flow cytometry

For analysis of F-actin content cells were labeled with Fixable Viability Dye eFluor™ 780 (Invitrogen™) and for the permeabilization was used 50 µL of Perm Buffer (Biolegend), 20 minutes at room temperature. The cells were washed with Perm Buffer, followed by staining with Alexa Fluor 647 phalloidin (Invitrogen™), and anti-WASp (F8; Santa Cruz Biotechnology Inc) conjugated to Zenon™ Human IgG Labeling Kit (Invitrogen™) in Perm Buffer for 30 minutes. After that the cells were washed two times, resuspended in FACS Buffer (DPBS supplemented with 5mM EDTA and 1% FSB) and assessed by flow cytometry.

### LFA-1 high affinity

0.5x10^6^ cells/mL were stimulated with ImmunoCult™ Human CD3/CD28/CD2 T Cell Activator (STEMCELL Technologies) or with anti-CD3 (OKT3, Biolegend) coated 96-well plate and incubated for 10, 30, and 60 minutes at 37 °C. The cells were stained with m24 for high affinity LFA-1 and TS2/4 for total anti-CD11a (both from Biolegend), followed by fixation with 2% PFA (Thermo Scientific®), and analyzed by flow cytometry.

### Statistical analysis

Prism software (GraphPad Prims 10) was used for statistical analysis. The results were analyzed by One-way ANOVA, or Two-way ANOVA as indicated.

## Results

### Recombinantly expressed WASp with missense variants reveals changes in protein conformation

To perform a systematic analysis of *WAS* variants, we selected 11 missense variants: two classified as WAS class I^4^ (E31K, G40R)^17–21^, two as WAS class II (A236G, D485N)^20,22–24^, six predicted as GOF (R268W, L270P, F271S, S272P, I290T, I294T)^13^, and one VUS from a patient with neutropenia clinically suspected to have XLN (R431W). We generated an AlphaFold 3.0^14^ prediction of wildtype (WT) WASp to visualize the location of the 11 missense variants (**Figure 1A**). E31K and G40R are located in the N-terminal WIP binding WH1 domain (residues 34–138)^25^. Variant A236G is in the disordered area and could disrupt the hydrophobic interactions in the Cdc42-WASp interaction interphase^26^. R268W, L270P, F271S, S272P, I290T, I294T^13^ are located in the WASp GBD domain (230-288)^27^, and R431W and D485N in the VCA region^7^ . We evaluated change in folding free energy compared to the WASp WT (ΔΔG=ΔG_WT_-ΔG_VARIANT_) using the DDmut software^15^. WAS variants L270P, F271S, I290T, and I294T were predicted to have a destabilizing effect on WASp structure, whereas the other seven variants did not change the folding free energy (**Figure 1B**). For biochemical analysis, we replaced the WH1 domain with an engineered spider silk tag (NT*) that enhances expression of highly disordered proteins for recombinant protein expression^28^, while preserving regions required for Cdc42 binding^27^ and WASp interactions with G-actin and the Arp2/3 complex^7^. This strategy enabled expression and purification of WT WASp and WASp variants under non-denaturing conditions for biochemical assays. We used native mass spectrometry of intact WASp to experimentally assess whether the *WAS* gene variants affected the conformational landscape of the protein^29^. Briefly, an unfolded protein with a large surface area can acquire a high number of charges and a broad charge state distribution in ESI-MS, whereas compact or globular proteins have narrow distributions with a low number of charges. When comparing mass spectra of all variants, we found that WASp WT, A236G, R268W, R431W, and D485N all obtained lower average charges, indicating globular or compact conformations (**Figure 1C-D**). L270P, F271S, S272P, I290T and I294T XLN variants, on the other hand, acquired significantly higher number of charges, indicating that these proteins preferentially occupy partially unfolded or extended states (**Figure 1C-D**). Molecular prediction combined with successful expression of recombinant WASp variants allowed us to define that five XLN variants preferentially adopted extended conformations.

**Figure 1.**
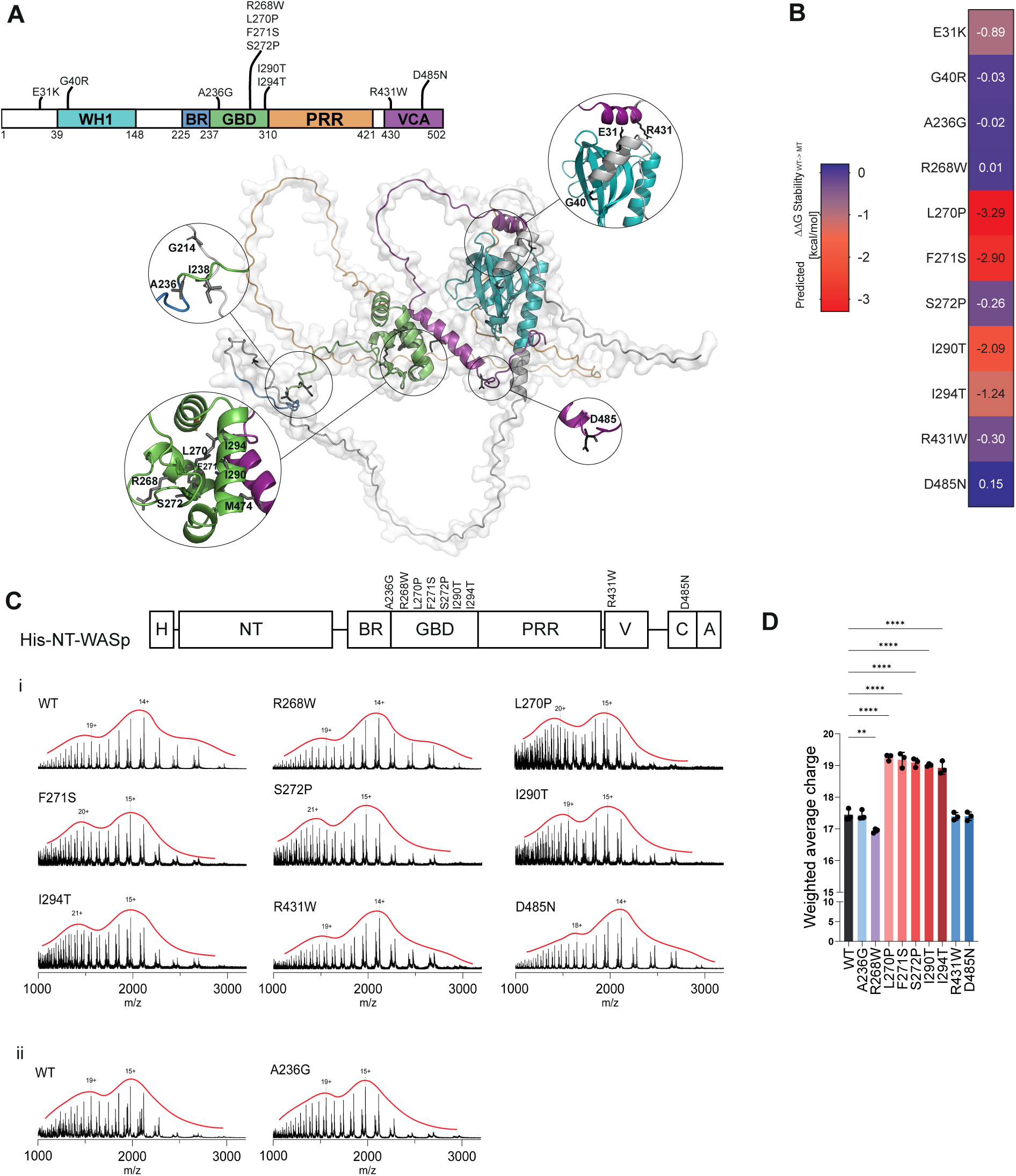
Distinctive effects on WASp stability of patient variants across the WASp structure. (A) Domain map of WASp with missense variants and WASp structure from AlphaFold 3.0. Close-up view of four regions where the variants are located. The amino acids that are substituted are shown as sticks. (B) DDG values upon amino acid substitution per variant where negative values indicate destabilization and positive values indicate stabilization. Purple indicates no changes in stabilization and red indicates decrease of stability. (C) His-NT*-WASp construct with indicated amino acid substitutions. Representative Native mass spectrum of each construct. (D) Quantification of the weighted average charge state. One-way ANOVA statistical test was performed. **P<0.002 *** P<0.0002**** P<0.0001.

### XLN variants of WASp show high rate of actin polymerization in the absence of Cdc42

To investigate the functional effects of WASp variants, we performed pyrene actin polymerization assays. WASp was mixed with purified Cdc42 loaded with GTP, pyrene actin at a ratio of 1:10 with non-labeled G-actin, and recombinant Arp2/3 complex in the presence of magnesium to stabilize newly formed actin filaments ^7,27,30,31^. When compared to G-actin, G-actin+Arp2/3, and G-actin+Arp2/3+Cdc42, the fluorescence intensity curve for WASp WT was two-fold higher indicated as time to reach half of maximal fluorescence in seconds (t_1/2_, **Figure 2A)**. The VCA domain alone showed rapid actin polymerization (**Figure 2A**). The WAS class II variants A236G^22^, D485N^19,32^, VUS R431W, and R268W had similar actin polymerization rate to WT WASp (**Figure 2A**). This analysis suggests that WASp with *WAS* variants was stable and active in the recruitment of free actin monomers and interaction with the Arp2/3 complex to induce pyrene actin polymerization. We next examined the XLN variants (L270P, F271S, S272P, I290T, I294T) and, similarly to VCA alone, they induced rapid rate of actin polymerization, showing that the VCA domain was fully exposed in the WASp with XLN variants (**Figure 2B**). We evaluated whether WAS variants could induce pyrene actin polymerization independently of Cdc42. WAS variants and VUS (A236G, R268W, R431W, D485N) led to actin polymerization rate similar to actin+Arp2/3 and G-actin alone, suggesting an autoinhibited WASp conformation with the VCA domain hidden (**Figure 2C**). In contrast, XLN variants exhibited robust actin polymerization in the absence of Cdc42, almost reaching the rate observed with VCA alone. (**Figure 2D**). The addition of Cdc42 further enhanced actin polymerization induced by the XLN variants, suggesting that the XLN variants did not completely disrupt WASp autoinhibited conformation. Of note, the disease-causing WAS variants A236G and D485N did not interfere with recombinant expression of NT*-tagged WASp variants, the molecular folding of WASp (**Figure 1**), nor the actin polymerization capacity in the presence of Cdc42 (**Figure 2**). This suggested that the cause of WAS disease in these patients may be related to the cellular stability of WASp.

**Figure 2.**
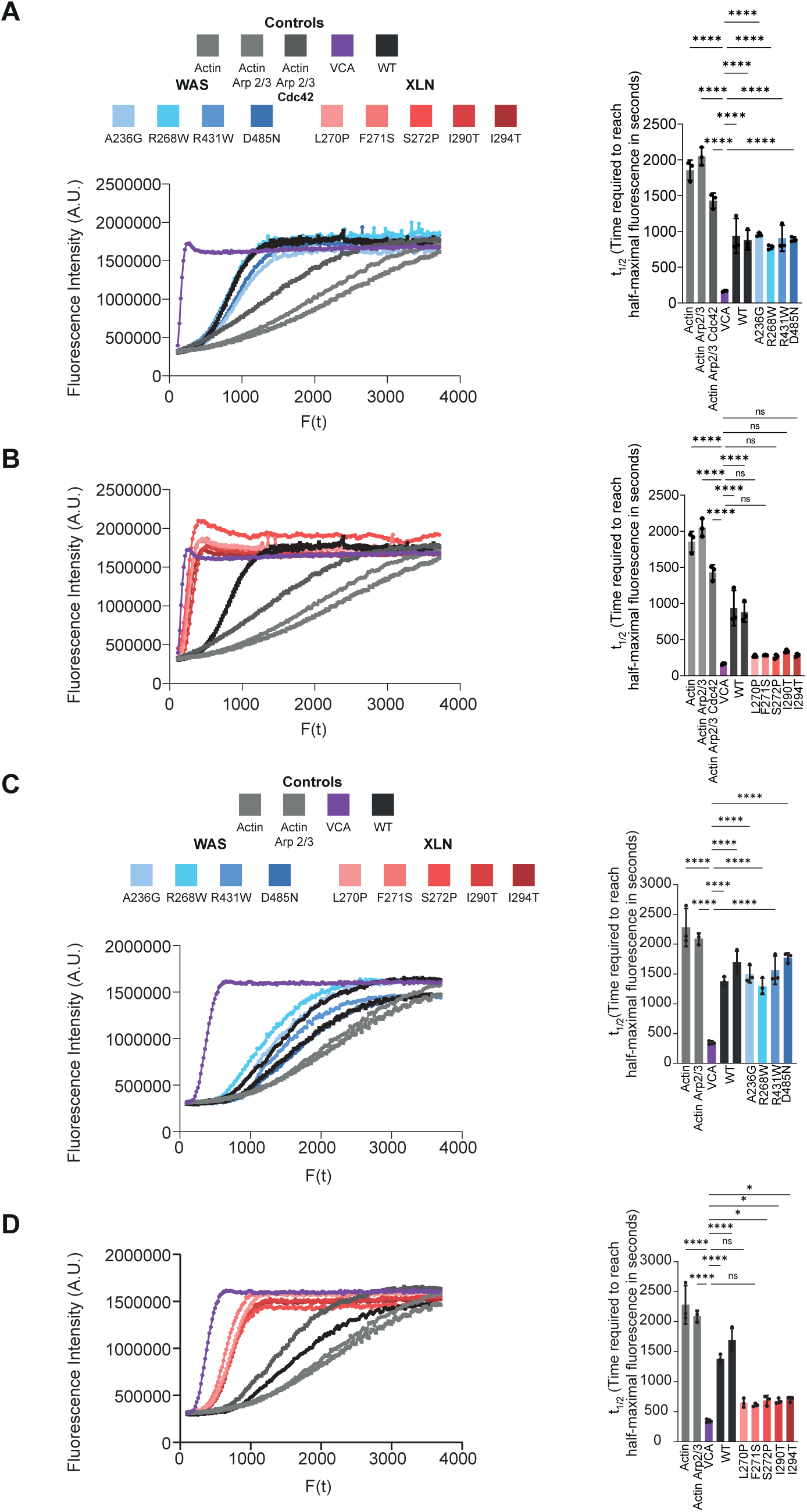
XLN variants lead to Cdc42 independent actin polymerization. The intensity of pyrene actin fluorescence reflected newly polymerized actin and was measured over time. (A and B) Representative graphs of pyrene actin polymerization time course of different WAS (A) and XLN (B) variants in presence of Cdc42-GTPgS, Arp2/3, pyrene actin at 1:10 with unlabeled actin, and magnesium. The time at which half of the fluorescence intensity occurred (*t* _1/2_) is indicated in the right panel (*n* = 3). (C) Actin polymerization without the WASp activator, Cdc42 of different WAS (C) and XLN (D) variants. Actin polymerization rate at *t* _1/2_ calculated from the polymerization curves without Cdc42 is indicated in the right panel (*n* = 3). One-way ANOVA statistical test was performed. **P<0.002 *** P<0.0002**** P<0.0001.

### CD4^+^ T cells harboring WAS variants have lower or absent WASp and reduced proliferation

To define the cellular consequence of WAS and XLN variants in a systematic comparison, we set up assays that can be performed on CD4^+^ T cells isolated from patient PBMCs. We focused on CD4^+^ T cells as they have been extensively studied in WAS patients ^33–36^, and less studied in XLN patients^13,37,38^. We included prospectively collected patient-derived CD4⁺ T cells harboring two WAS variants (E31K and G40R), two predicted XLN variants (F271S, I294T)^13^, and two VUS (R268W, R431W). Patient-derived cells were compared with healthy donors (HD) cells isolated from two unrelated donors (HD1 and HD4), the sister of the R268W patient (HD2), the mothers of patients carrying G40R, R86C, and R439W variants (HD3, HD5, and HD6, respectively). CD3^+^CD4^+^ T cells were FACS sorted from patient-derived PBMCs (**Supplemental Figure 1A)** and stimulated with anti-CD2/CD3/CD28 for 10 days. All cells were subsequently cryopreserved and later thawed for analysis (**Figure 3A**). WASp expression was absent in E31K CD4⁺ T cells and reduced in G40R CD4⁺ T cells compared with healthy donor and XLN CD4⁺ T cells. (**Figure 3B**). Upon anti-CD3 activation, the E31K and G40R CD4+ T cells had reduced proliferation when compared to HD cells, suggesting that reduced WASp expression in G40R cells was insufficient to induce CD4⁺ T cell proliferation following anti-CD3 activation. No significant differences in CD3 expression were detected across the samples (**Supplemental Figure 1B).** The XLN (F271S, I294T) CD4^+^ T cells proliferated efficiently and obtained 1-2 additional cell divisions when compared to HD cells (**Figure 3C**). The VUS R268W CD4^+^ T cells proliferated comparably to HD cells. The proliferative response of G40R CD4^+^ T cells was rescued with optimal T cell activation using anti-CD2/CD3/CD28 stimulation and reached similar proliferation to HD and XLN CD4^+^ T cells (**Figure 3D**). However, E31K CD4^+^ T cells with absent WASp had lower proliferation rate compared to the HD cells also in response to anti-CD2/CD3/CD28 stimulation (**Figure 3D**). This data suggest that presence of WASp is required for proliferation upon robust CD4^+^ T cell activation. Moreover, increased WASp activity in XLN cells induced higher proliferative capacity of CD4^+^ T cells at low T cell activation by anti-CD3 stimulation only.

**Figure 3.**
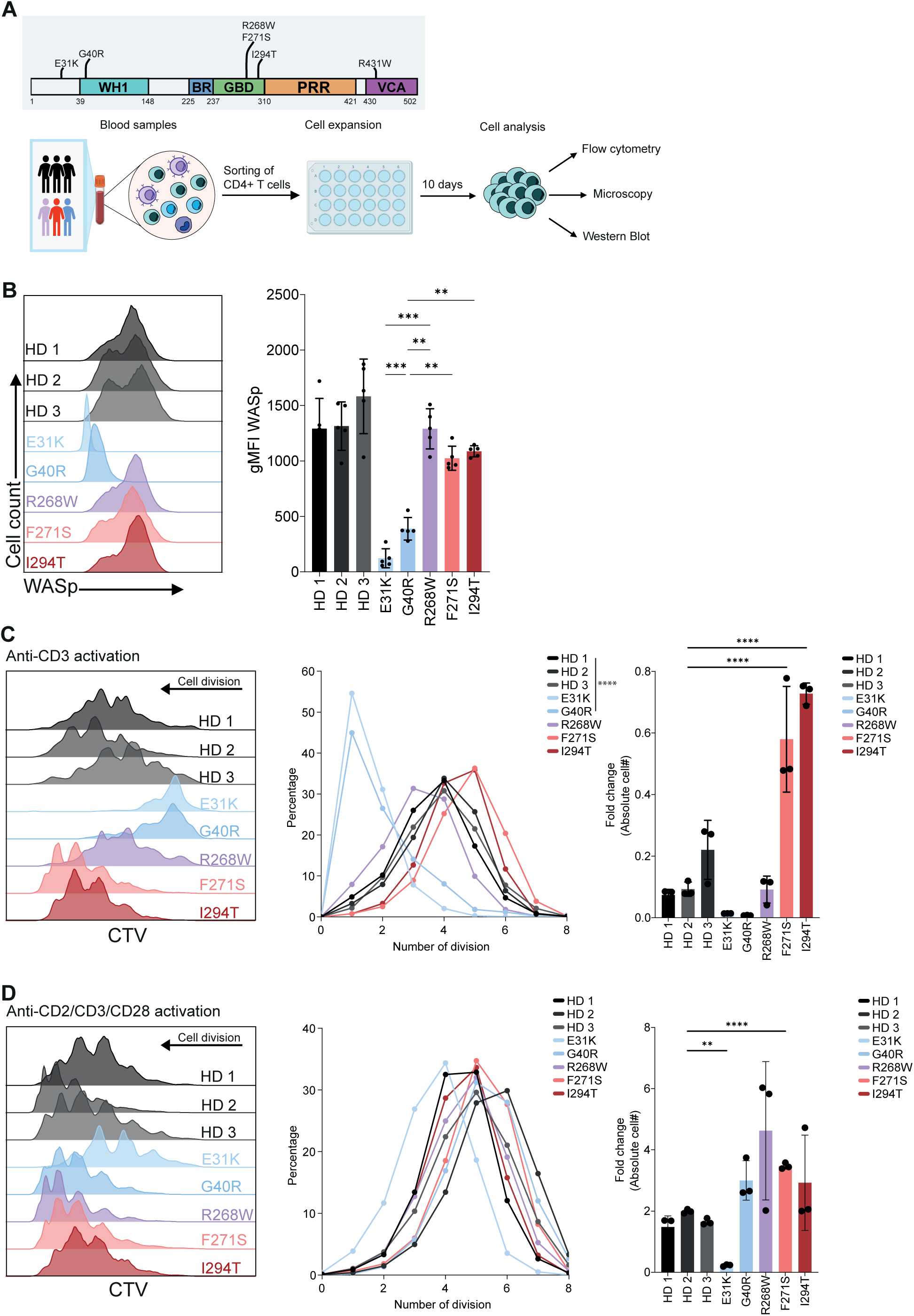
WAS CD4^+^ T cells have poor proliferation and XLN CD4^+^ T cell enhanced proliferation upon anti-CD3 stimulation. (A) Distribution of patient *WAS* gene variants across the WASp domains. Schematic diagram of the methods used. (B) Representative flow cytometry histogram plots for WASp. Quantification of gMFI WASp in different samples (n=5). (C-D) Representative histogram of proliferation of CD4^+^ T cells in patient samples and healthy controls stained with CellTrace Violet (CTV) after 5 days of anti-CD3 (C) or anti-CD2/CD3/CD28 (D) stimulation. (Middle panel) Percentage of proliferating T cells according to number of divisions as estimate by CTV dilution. (Right panel) Fold increase of number of cells after 5 days stimulation (n=3). Statistical significance was determined by One-way ANOVA. **P<0.002 *** P<0.0002****P<0.0001.

### WASp expression level correlates with F-actin content

To assess the functional activity of WASp *in vitro*, we measured polymerized actin using phalloidin. E31K cells with absent WASp had reduced F-actin content, whereas G40R cells showed levels comparable to healthy donors, as well as the VUS R268W cells. The XLN I294T cells had increased F-actin content (**Figure 4A**). To understand the relationship between WASp level and F-actin content, we performed correlation analysis. We observed a positive correlation between WASp expression and F-actin content, best demonstrated in WAS (E31K, G40R) variants *versus* XLN (F271S, I294T) variants (**Figure 4B**). The R268W VUS were similar to HD cells in terms of WASp and F-actin content. As a functional read out of F-actin content, we examined the capacity of CD4^+^ T cells to spread on anti-CD3 antibody-coated cover slips by confocal microscopy. WAS (E31K, G40R) cells had reduced F-actin content when compared to age matched HD cells (HD2), whereas XLN cells (F271S, I294T) had higher F-actin content when compared to HD cells (**Figure 4C, Supplemental Figure 2A**). This data suggests that in addition to WASp expression by flow cytometry, analysis of F-actin by flow cytometry and microscopy is helpful in categorizing *WAS* gene variants.

**Figure 4.**
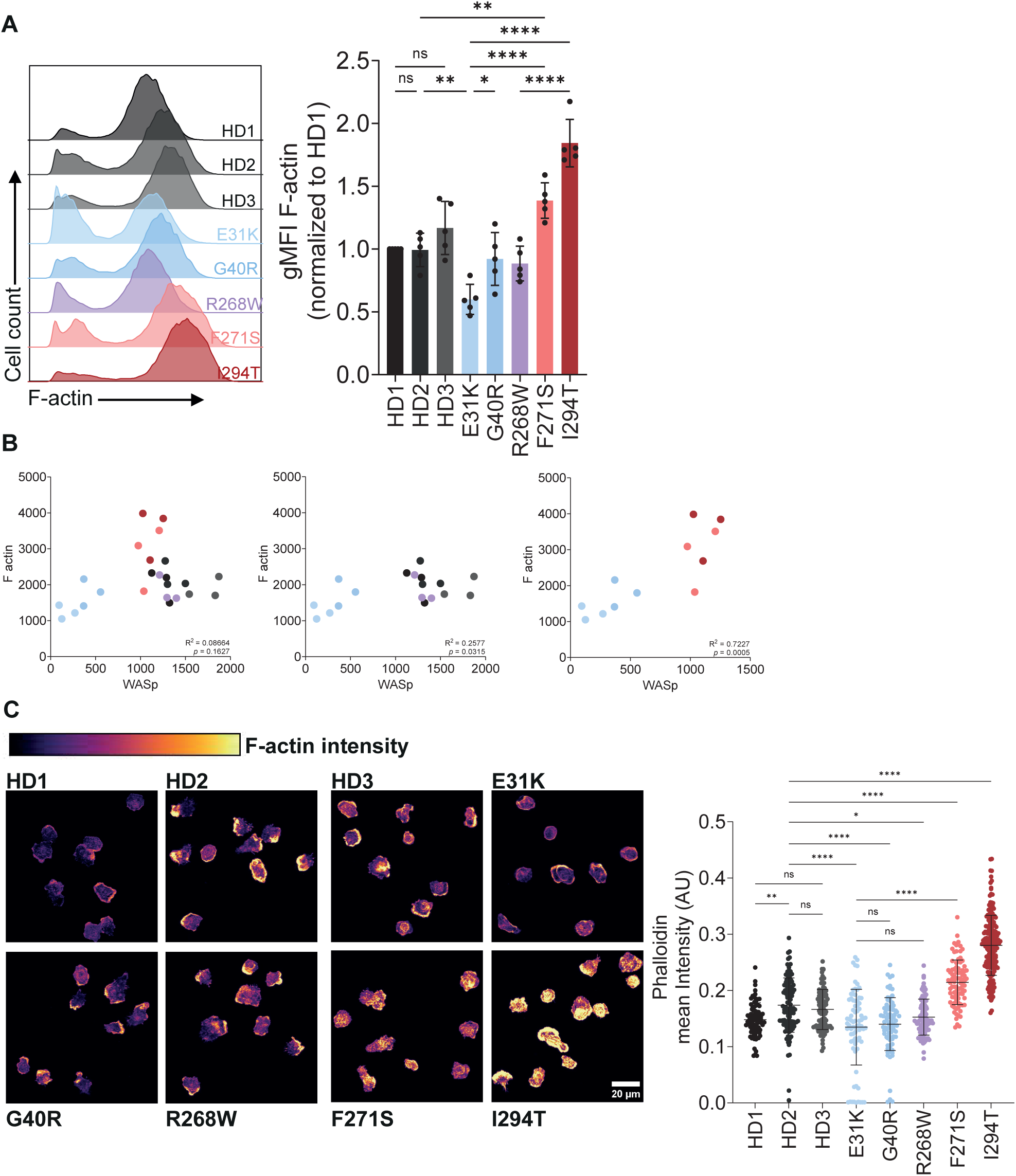
F-actin content is decreased in WAS CD4^+^ T cells and increased in XLN CD4^+^ T cells. (A) Representative flow cytometry histrogram for F-actin content in CD4^+^ T cells. (Right panel) Quantification of gMFI F-actin normalized by HD1 F-actin content (n=5). (B) Pearson’s correlation between F-actin and WASp intensities comparing all samples, WAS and HD samples, and WAS and XLN samples. (C) Representative confocal fluorescence microscopy images of CD4^+^ T cells from patients on anti-CD3 coated glass. (Left panel). Quantification of F-actin content by phalloidin intensity (Right panel). Data shown from one independent experiment conducted on primary CD4^+^ T cells. In total at least 70 cells were studied. Statistical significance was determined by One-way ANOVA. **P<0.002 *** P<0.0002****P<0.0001.

### *WAS* gene variants affect TCR induced phosphorylation of WASp tyrosine-291 and activation of high affinity LFA-1

To assess the effects of WAS and XLN variants on WASp activation in CD4^+^ T cells, phosphorylation of tyrosine-291 (Y291) was examined, which becomes exposed upon Cdc42 binding and release of WASp autoinhibition, thus revealing the active, open conformation of WASp. Western blotting analysis confirmed undetectable WASp expression in E31K cells and reduced WASp expression in G40R cells (**Figure 5A**). A phosphorylation-specific Y291 WASp antibody did not detect WASp in WAS and HD cells, but detected pY291-WASp in XLN (F271S, I294T) cells (**Figure 5A**), demonstrating an open and active conformation of XLN variants. To enrich WASp and thereby pY291-WASp, we immunoprecipitated WASp from CD4^+^ T cell lysates followed by detection of pY291-WASp. XLN (I294T) cells had increased pY291-WASp/total WASp ratio when compared to HD cells, whereas WAS (G40R), with low expression of WASp, had reduced pY291-WASp/total WASp ratio (**Figure 5B**). The VUS R268W and R431W cells had similar pY291-WASp when compared to WASp WT cells (**Figure 5B**). This indicates that reduced WASp expression together with decreased capacity to disrupt the autoinhibited conformation of WASp may influence the cellular response of WAS G40R CD4^+^ T cells. Moreover, XLN (F271S, I294T) CD4^+^ T cells had increased baseline pY291-WASp similarly to neutrophils^39^ and B cells^13^ harboring murine XLN variants.

**Figure 5.**
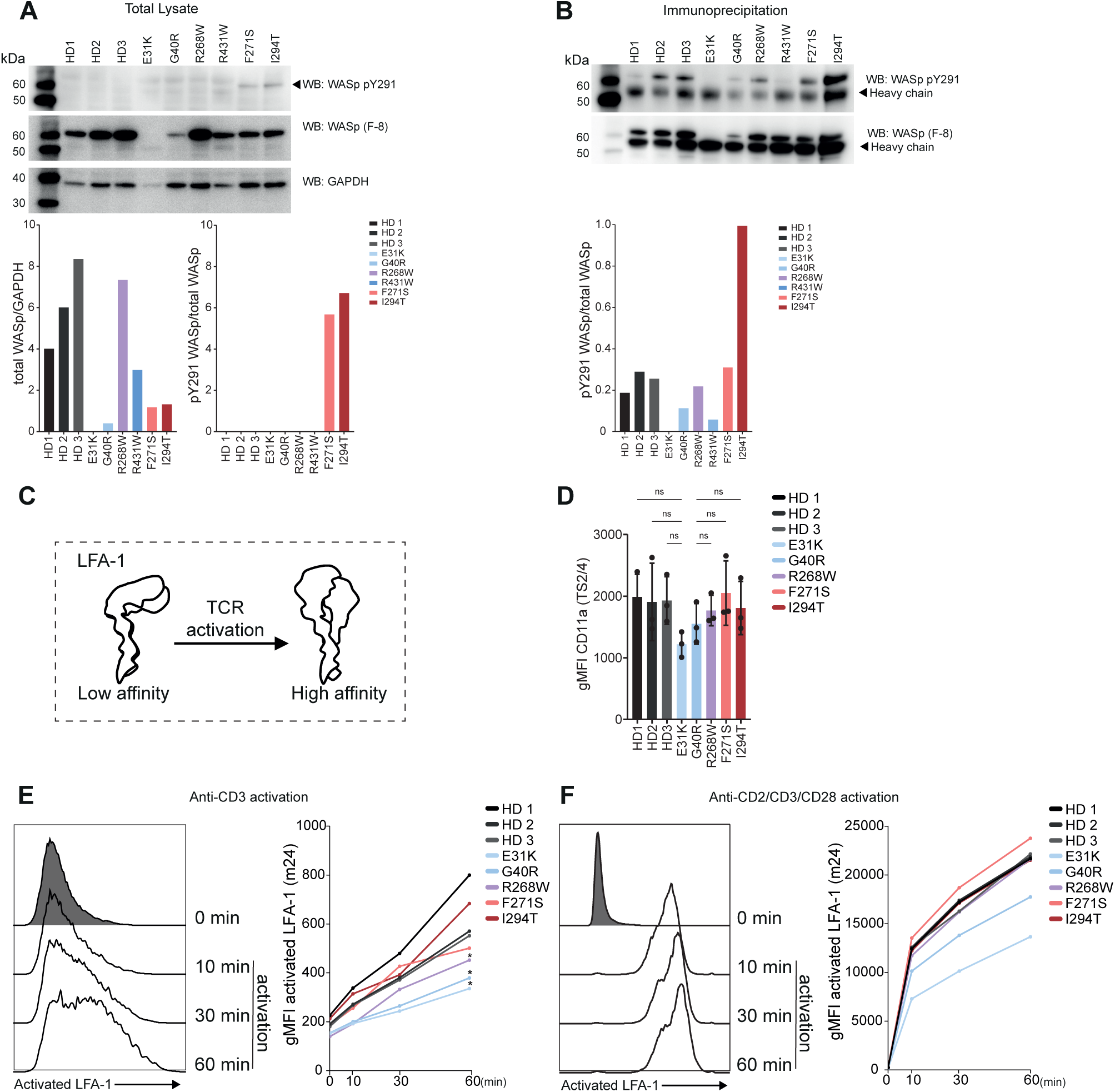
Phospho-Y291 of WASp and upregulation of active LFA-1 in WAS and XLN CD4^+^ T cells. (A) Western blot analysis of total CD4^+^ T cell protein lysates from indicated *WAS* variants and detection with anti-WASp (total) and anti-pY291-WASp. Bar chart indicates WASp/GAPDH ratio and ratio of pY291-WASp/total WASp of one experiment. The WAS patient values were not used for ratio evaluation because of none/low levels of WASp expression. (B) Immunoprecipitation of WASp from the total CD4^+^ T cell protein lysates and bar chart of ratio pY291 WASp/total WASp. (C) Scheme of LFA-1 activation after TCR engagement. (D) gMFI of total LFA-1 (antibody clone TS2/4) in CD4^+^ T cells by flow cytometry. Non-significant with one-way Anova. (E) Representative flow cytometry histograms of HD1 CD4^+^ T cells of active LFA-1 (antibody clone m24) overtime upon anti-CD3 stimulation (1 representative assay of 3). (Right panel) gMFI of high affinity LFA-1 at different time points (0, 10, 30 and 60 min). (F) Representative flow cytometry histograms of HD1 CD4^+^ T cells of active LFA-1 overtime upon anti-CD2/CD3/CD28 activation (1 representative assay of 3). Right panel) gMFI of high affinity LFA-1 at different time points (0, 10, 30 and 60 min). Data presented as the mean from 3 separate experiments (D-F). *Denotes significant increase of active LFA-1 compared with HD1 (p < 0.03) using Two-way ANOVA.

We examined how *WAS* variants may impact on upregulation of high affinity LFA-1 upon TCR activation by anti-CD3 and anti-CD2/CD3/CD28 stimulation (**Figure 5C**). CD4^+^ T cells with WAS (E31K, G40R), XLN (F271S, I294T), and VUS R268W variants had similar expression of total LFA-1 (CD11a) (**Figure 5D**). Upon anti-CD3 activation, WAS (E31K, G40R), and VUS R268W cells had reduced capacity to upregulate active LFA-1 when compared to HD cells (**Figure 5E, Supplemental Figure 3C-D**). XLN (F271S, I294T) CD4^+^ T cells upregulated high affinity LFA-1 similarly to HD cells. In response to stronger anti-CD2/CD3/CD28 activation, WAS cells (E31K, G40R) upregulated active LFA-1, albeit less than HD, VUS R268W, and XLN (F271S, I294T) cells (**Figure 5F and Supplemental Figure 3C-D**). Together, this data shows that analysis of pY291-WASp and upregulation of active LFA-1 help categorizing *WAS* gene variants into LOF or GOF. VUS R268W, R431W may have only moderate effect on WASp activity and behave largely as WASp WT in HD CD4^+^ T cells.

### A set of biochemical and cellular assays provides a molecular map of *WAS* gene variants

A heat map generated in GraphPad Prism was used to integrate in silico, biochemical, and cellular data across WAS variants, with color gradients indicating functional effects ranging from gain-of-function (red) to loss-of-function (blue). (**Figure 6A**). Differentiation between WAS-associated variants and VUS remains challenging. AlphaFold 3.0 modeling was applied to investigate functional interpretation of VCA variants in WAS class II, revealing <3.5 Å contacts between WASp-D485 and Arp3-H371 residues in the interaction interphase (**Supplemental Figure 4**). Our functional analysis of CD4^+^ T cells indicated that combined analysis of WASp and F-actin by flow cytometry may help stratify *WAS* gene variants. We validated this approach on new prospectively-collected patients with typical WAS thrombocytopenia, carrying two variants previously reported in WAS (T45M, R86C) ^17,18,20,23,40–42^ and four VUS (G214A, I238T, R439W, M474T). We detected very low to absent WASp expression in T45M, R86C patient CD4^+^ T cell from PBMCs and in G214A CD4^+^ T cell blasts (**Figure 6B**, left panel). WAS T45M and G214A had lower F-actin content when compared to HD PBMCs, whereas R86C had F-actin similarly to the HD cells (**Figure 6B**). VUS I238T, R439W, and M474T had normal expression of WASp, but reduced F-actin, suggesting normal expression of a dysfunctional WASp. The biochemical and cellular assays developed here should be combined with long term follow up, especially for VUS that are difficult to distinguish from WASp WT. Correlation analysis of WASp expression and F-actin content provided a clear stratification of *WAS* gene variants with low/absent WASp and low/absent F-actin (**Figure 6C**). In cells harboring *WAS* gene variants leading to normal expression of WASp, reduced F-actin may indicate protein dysfunction.

**Figure 6.**
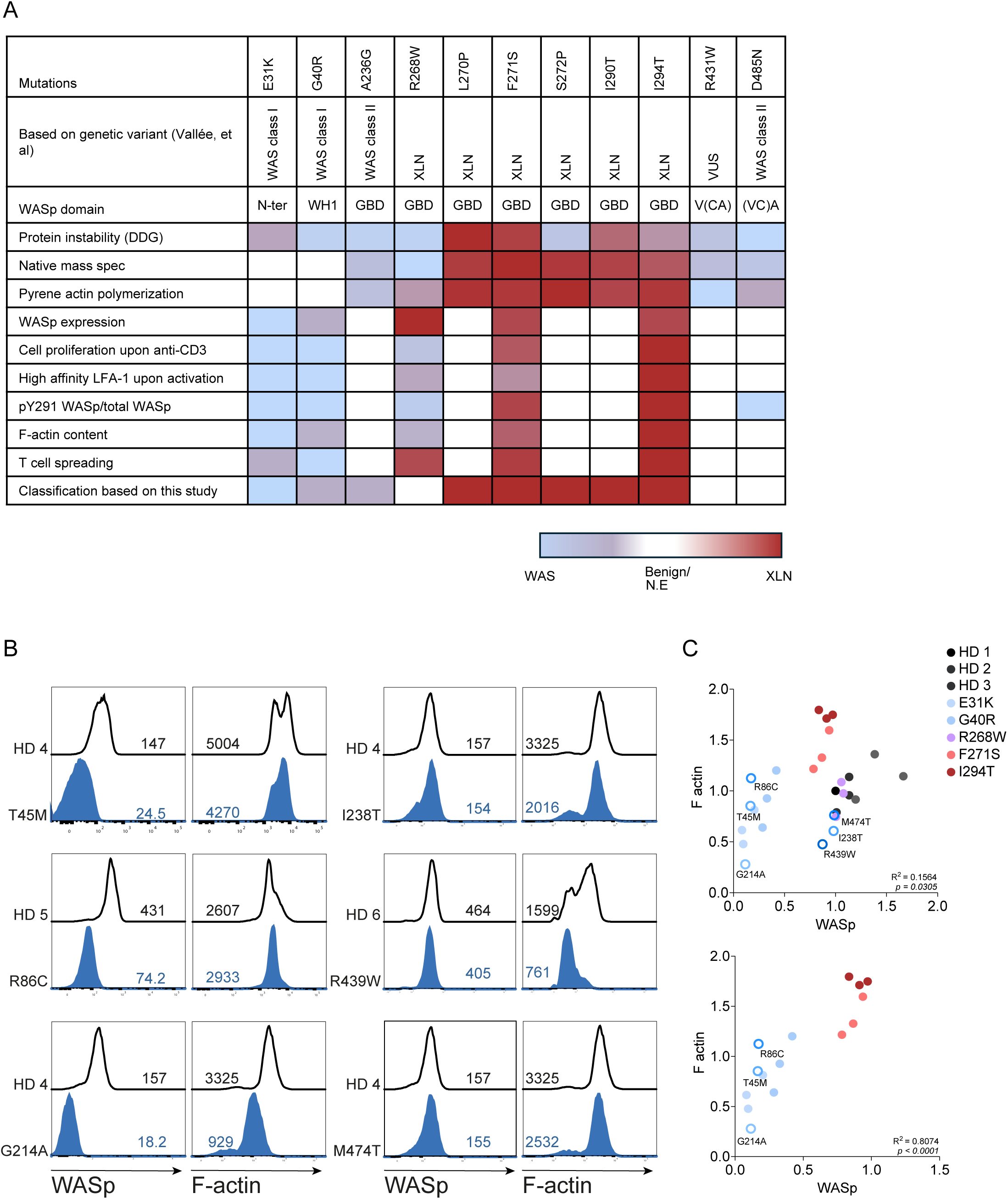
Molecular and cellular map of 11 WAS variants. (A) Effects of each *WAS* gene variant and predicted disease severity. A heat map was generated in GraphPad Prism according to the results obtained. Blue indicates a low response, white indicates mild or absent response, and red denotes a high response for the selected parameter. White rectangle can also correspond to variants not evaluated in the specified assay. Reference to published work is indicated in parentheses. (B) Patient CD4^+^ lymphocyte staining for WASp and F-actin compared to Healthy Donor samples. (C) Correlations between F-actin content and WASp staining of all samples (upper panel) and only WAS and XLN samples (lower panel). Data was analyzed by Pearson’s correlation.

## Discussion

*WAS* gene variants may differently affect WASp stability and conformation, as well as actin polymerization capacity and consequently, the function of hematopoietic cells. WASp expression by flow cytometry is widely used to establish the diagnosis of WAS, however, it can be normal in some cases and has demonstrated insufficient correlation with severity of the disease^43^. For XLN diagnosis, WASp expression is misleading as the protein may be prone to degradation due to the open conformation^39^, and functional assays are needed to determine higher WASp activity. We took a systematic approach to compare 11 *WAS* gene variants including four WAS, one VUS, and six predicted XLN variants. NT*-tag WASp allowed us to directly screen for differences in protein conformation and actin polymerization capacity. We developed a set of flow cytometry-based assays using CD4^+^ T cells that distinguished graded WASp expression and F-actin content, enabling discrimination among WAS, XLN, and likely benign *WAS* gene variants.

Due to the highly disordered structure of WASp, most biochemical data of WASp structural regulation come from mini-constructs of WASp and N-WASp, revealing that WASp/N-WASp has a closed conformation and binding to Cdc42 releases the autoinhibition and exposes the VCA domain^7–9^. We predicted how WASp variants affected the destabilization of WASp structure using AlphaFold 3.0^14^ and DDmut prediction tool^15^. The four XLN variants L270P, F271S, I290T, I294T led to reduced stability, whereas variants R268W, S272P and WAS variants had no predicted effect on stability on WASp. Compared with compact proteins, disordered proteins usually display a wider distribution of charge states upon ionization^29^. NT*-tagged WASp with XLN variants L270P, F271S, S272P, I290T, I294T showed an increased average charge state by native mass spectrometry, whereas WASp WT, R268W (previously predicted XLN), WAS variants (A236G, D485N), and VUS (R431W) had lower average charge state, suggesting a compact conformation. Using circular dichroism of a mini-WASp construct containing the GBD-VCA domain, the structures of L270P^37^ and I294T^44^ protein was shown to have a lower melting temperature than WT WASp, indicating an increased tendency to unfold. Together, these findings demonstrate that structural instability and unfolding are characteristic features of XLN variants of the *WAS* gene.

We selected two WAS missense variants (A236G and D485N) known to be associated with reduced WASp expression ^20,22–24^. A236G is in the N-terminal part of the GBD, while D485N lies near serine residues 483 and 484, whose phosphorylation by casein kinase 2 induces a 2-fold increase in pyrene actin polymerization^45^. It remains unclear how WASp missense variants outside the WH1 domain lead to WAS. We predicted that the A236G variant may affect Cdc42 binding, whereas the D485N variant may lead to altered binding to Arp2/3. However, WAS A236G and D485N had pyrene actin polymerization rates comparable to WASp WT, suggesting that the variants did not disrupt the interaction with Cdc42, Arp2/3, or globular actin, at least not when in excess *in vitro*. Patient cells with A236G or D485N variants were not available, these variants may affect WASp cellular function in ways not modelled by the pyrene actin assay. Yeasts lacking the WASp ortholog, Las17, have impaired growth. Human WASp A236G rescues this defect, as a readout for WASp activity, whereas D485N led to reduced cell growth^46^. The WAS I481N variant, associated with intermittent thrombocytopenia and persistently small platelets without other major clinical WAS manifestations, leads to normal WASp expression^47^ and similarly complements growth in Las17-deficient yeast^46^. Our biochemical data of WAS A236G and D485N showed normal *in vitro* actin polymerization, but D485N led to reduced cellular function^46^, and both lead to severe clinical disease ^20,22–24^. This has implication for disease severity prediction in patients with residual WASp expression as increasing expression of WASp variant(s) may be insufficient to fully reconstitute protein function in vivo.

We set up functional assays that allowed for stratification of WAS, XLN, and benign variants, and that can be easily adoptable to examine CD4^+^ T cells from a small volume of patient blood sample. WAS class I variants affecting the WH1 domain (E31K, G40R) showed reduced or absent WASp expression and activity, resulting in severe functional impairment when WASp was entirely absent. In patients with WAS class I variants, overall and event-free survival were not influenced by the presence or absence of WASp expression^19^. However, compared with patients with WAS class II variants, they have milder course of the disease or delayed onset of severe disease manifestations^4^. In our study, the partial recovery of proliferation observed in WAS CD4^+^ T cells after full activation suggests that robust T cell stimulation can, to some extent, bypass the requirement for WASp. This agrees with earlier studies demonstrating that reduced proliferation of murine WASp null T cells stimulated with anti-CD3 can be partly recovered with addition of IL-2^48–52^. Murine WASp null T cells have defects in actin cytoskeletal dynamics following TCR activation^36,52,53^. Consistent with this, we observed reduced F-actin content in WASp-null CD4^+^ T cells (E31K), whereas XLN variant I294T displayed increased F-actin levels, in agreement with murine XLN T cells^54^.

Upon TCR engagement, actin cytoskeleton remodeling is required for productive T cell activation^55^. Integrin mediated adhesion is required to stabilize the immune synapse^56^. WASp supports LFA-1 activation and nanocluster organization, both essential for CD8^+^ T cell cytotoxic activity^57^. Following gene therapy for WAS, immune synapse formation including LFA-1 activation was restored^58^. We observed reduced upregulation of high affinity LFA-1 in WAS E31K and G40R CD4^+^ T cells, associated with decreased F-actin content in E31K CD4^+^ T cells. By comparing CD4^+^ T cells lacking WASp expression with XLN cells, we observed a clear association between WASp Y291 phosphorylation and LFA-1 activation.

The systematic approach to analyze *WAS* gene variants expands the clinical spectrum of WASp related diseases and highlights the importance of determining biochemical and cellular function of *WAS* gene variants, especially for WAS variants outside exon 1 and 2 and for VUS variants. Our data delineates how loss-of-function and gain-of-function variants in WASp differentially affect its structural stability, actin polymerization capacity, and CD4^+^ T cell function.

## Acknowledgement

We are grateful to the patients and their families for providing blood samples.

This work was supported by postdoctoral fellowships from Wenner-Gren Foundations to L.G.P., from Cancer Society to M.K. and J.R., a CAPES-STINT joint grant to L.G.P., M.R.B, G.P.F. and L.S.W., the Swedish Research Council and Cancer Society to L.S.W., and Worldwide Cancer Research, Childhood Cancer Fund, Radiumhemmet Research Funds, and Karolinska Institutet to L.S.W. L.S.W. holds a senior research position awarded by the Childhood Cancer fund.

## Author Contribution

L.G.P., M.R.B., A.L., M.L. and L.S.W. conceptualized the study and designed the research, L.G.P., M.R.B., A.L., R.C.V., K.R., J.S., M.H., M.K., and J.R. performed the experiments and analyzed the data, U.T., F.K., M.S., A.vdV., J.B., K.R., A.P., A.S., O.E., G.P.F., and M.L. contributed with patient samples, critical tools, and discussions, L.G.P. and L.S.W. wrote the manuscript, and all authors edited the manuscript.

